# LMO7 deficiency reveals the significance of the cuticular plate for hearing function

**DOI:** 10.1101/334052

**Authors:** Ting-Ting Du, James B. Dewey, Elizabeth L. Wagner, Shimon P. Francis, Edward Perez-Reyes, Wenhao Xu, John S Oghalai, Jung-Bum Shin

**Affiliations:** Department of Neuroscience, University of Virginia, Charlottesville, Virginia, USA; Caruso Department of Otolaryngology-Head and Neck Surgery, University of Southern California, Los Angeles, California, USA; Genetically Engineered Murine Model (GEMM) core, University of Virginia, Charlottesville, Virginia, USA; Department of Pharmacology, University of Virginia, Charlottesville, Virginia, USA

## Abstract

Sensory hair cells, the mechanoreceptors of the auditory and vestibular system, harbor two specialized organelles, the hair bundle and the cuticular plate. Both subcellular structures have adapted to facilitate the remarkable sensitivity and speed of hair cell mechanotransduction. While the mechanosensory hair bundle is extensively studied, the molecules and mechanisms mediating the development and function of the cuticular plate are poorly understood. The cuticular plate is believed to provide a rigid foundation for stereociliar pivot movements, but specifics about its function, especially the significance of its integrity for long-term maintenance of hair cell mechanotransduction, are not known. In this study, we describe the discovery of a hair cell protein called LIM only protein 7 (LMO7). In the hair cell, LMO7 is specifically localized in the cuticular plate. *Lmo7 KO* mice suffer multiple deficiencies in the cuticular plate, including reduced filamentous actin density and abnormal length and distribution of stereociliar rootlets. In addition to the cuticular plate defects, older *Lmo7 KO* mice develop abnormalities in inner hair cell stereocilia. Together, these defects affect cochlear tuning and sensitivity and give rise to late-onset progressive hearing loss.

## Introduction

In few other cell types is the principle of “Form follows Function” as evident as in the sensory hair cell. The hair cell’s subcellular structures are optimally designed to facilitate hair cell mechanotransduction, the process by which mechanical energy from sound and head movements are converted into cellular receptor potentials. Two specialized structures in the hair cell, the hair bundle and the cuticular plate, are essential for hair cell mechanotransduction ^1-7^. Both are hair cell-specific elaborations of structures found in other microvilli-bearing cells, such as the intestinal brush border cells ^8,9^. The hair bundle, an array of microvilli arranged in a staircase-like fashion, harbors the mechanotransduction complex. A substantial body of research has identified the mechanisms essential for the morphogenesis and function of the hair bundle ^10-16^. In contrast, the molecular composition and significance of the cuticular plate, a structure analogous to the brush border cell’s terminal web, is poorly understood. The cuticular plate is believed to provide a mechanical foundation for the stereocilia, which are inserted into the cuticular plate ^17-20^. A stiff stereociliar insertion point ensures that vibration energy is fully converted into stereocilia pivot movement, and not diminished by non-productive cuticular plate deformations. This notion is supported by electron microscopy-based ultrastructural studies, which demonstrate that the cuticular plate is reinforced by a dense network of actin filaments, crosslinked by actin-binding proteins such as spectrin ^21^. In addition to providing a mechanical foundation, the cuticular plate is also believed to be involved in selective apical trafficking of proteins and vesicles ^22^. However, specifics about the function and formation of the cuticular plate, especially the significance of its integrity for long-term maintenance of hair cell function, are unknown. This gap in knowledge is in part attributable to the lack of molecular tools to manipulate the cuticular plate specifically. Molecular studies have uncovered a few resident proteins such as spectrin, tropomyosin, supervillin ^23-27^, but loss-of-function studies for these proteins have not been undertaken to date.

In this study, we report the discovery of a novel component of the cuticular plate, a hair cell-enriched protein called LIM only protein 7 (LMO7). LMO7 contains a calponin homology (CH) domain, a PDZ domain, and a LIM domain, and was reported to be involved in protein-protein interactions at adherens junctions and focal adhesions ^28,29^. LMO7 deficiencies were reported to increase susceptibility to spontaneous lung cancer and Emery-Dreifuss Muscular Dystrophy (EDMD) ^30,31^. In addition, LMO7 was revealed to play a role in the regulation of actin dynamics through the Rho-dependent MRTF-SRF signaling pathway ^32^. Our studies show that in the hair cell, LMO7 is specifically localized to the cuticular plate and intercellular junctions. In LMO7-deficient mice, hair cells exhibit reduced F-actin staining in the cuticular plate, along with abnormal distribution of stereocilia rootlets. In addition, older *Lmo7 KO* mice develop abnormalities in the stereocilia of inner hair cells. These morphological defects affect tuning and sensitivity of cochlear vibrations, and cause late-onset progressive hearing loss.

## Results

### LMO7 is an abundant component of the hair cell cuticular plate

We previously used a peptide mass spectrometry-based strategy to characterize the hair bundle proteome of chick vestibular organs, and identified LIM-only protein 7 (LMO7) as a potential hair bundle enriched protein ^33,34^ (**Fig. 1b**). Subsequent immunolocalization analysis showed that LMO7 is expressed in the mouse inner ear and highly enriched in the sensory hair cells. Contrary to our expectations, however, its localization was restricted to the cuticular plate and the intercellular junctions (**Fig. 1d-f**). The presence of LMO7 in the hair bundle preparation was likely due to co-purification of cuticular plate material during the hair bundle twist-off purification. This is evident when “hair bundle” preparations from mouse utricles, still embedded in the agarose matrix used for the twist-off method, are immunolabeled with LMO7-specific antibodies (**Fig. 1c**). LMO7 immunoreactivity is abundant in the cuticular plate, but not in the hair bundle.

**Figure 1.**
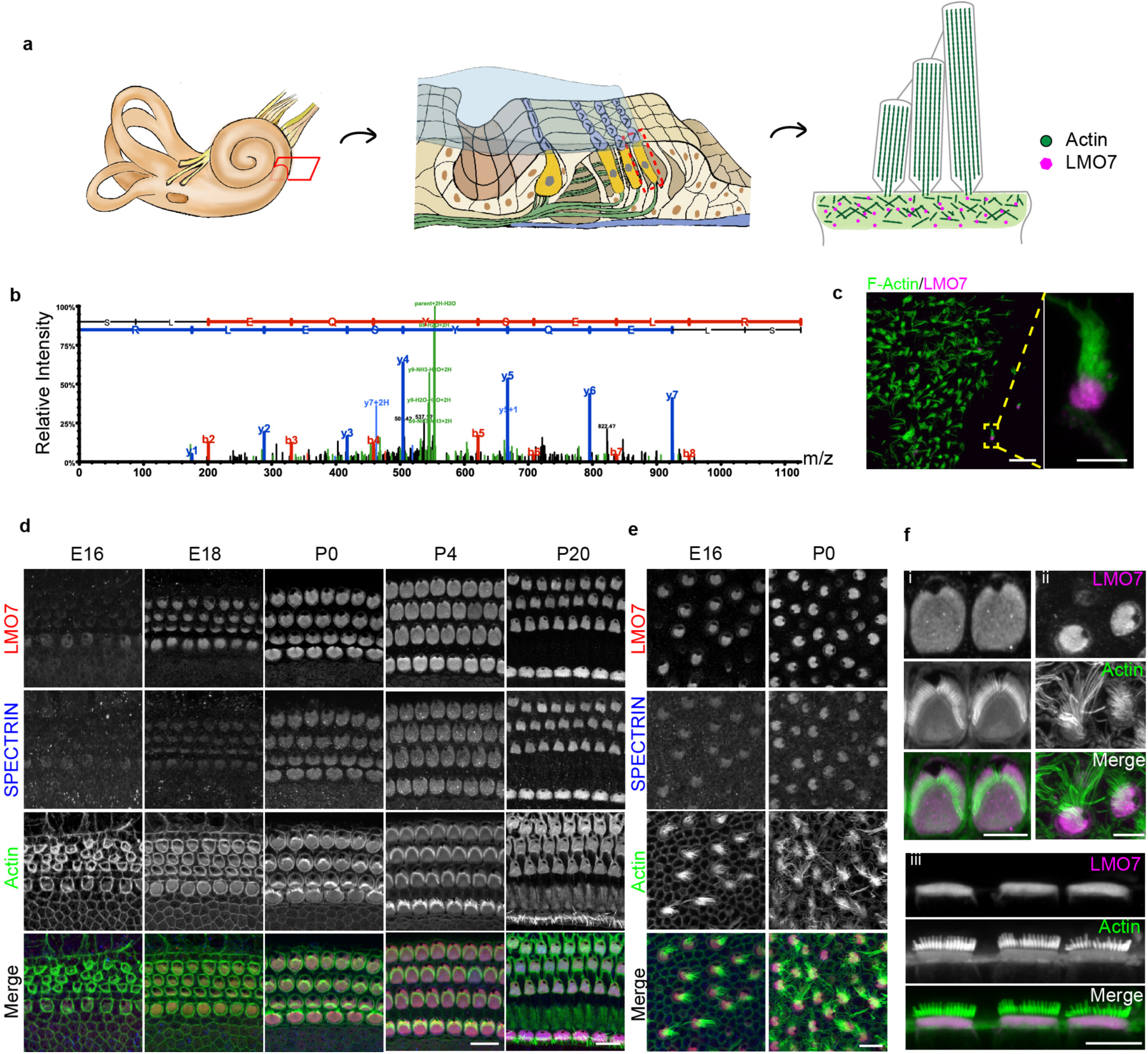
Lmo7 is a component of the hair cell cuticular plate and is specifically localized to the cuticular plate. **a.** Schematic representation of inner ear organization and the apical structures of the hair cell. **b**. MS/MS spectrum of a representative peptide of chick LMO7, identified by LC-MS/MS on isolated chick hair bundles. The data for the spectrum was obtained from a previously published dataset ^34^. **c**. LMO7 immunoreactivity (magenta) in isolated mouse hair bundles confirmed its presence in cuticular plate (labeled by phalloidin in green). Scale bars, 20 μm (overview), 5 μm (panel magnification). **d, e**. Immunocytochemical analysis of LMO7 expression in the mouse cochlea and utricle of various ages. LMO7 expression is detected in hair cells at E16, with the initial emergence of the hair bundle. Scale bar, 10 μm. **f**, (i,ii) Higher-magnification views of LMO7 immunoreactivity in the mouse cochlear and vestibular hair cell. LMO7 localization is restricted to the cuticular plate and the intercellular junctions. (iii) Side view of LMO7 expression in the inner hair cell. Scale bar, 5 μm.

We next examined the spatiotemporal expression of LMO7 in the mouse inner ear using immunohistochemistry at various stages of development. Initial LMO7 expression in the embryonic cochlea coincides with the emergence of hair bundles at embryonic day 16 (E16), and its expression increases and continues into mature stages (**Fig. 1d**). Likewise, LMO7 is abundant during development of the utricle cuticular plate (**Fig. 1e**). Overall, the spatiotemporal expression of LMO7 is similar to Spectrin which has been reported as an actin-binding protein localized in the cuticular plate (**Fig. 1d,e**). In summary, LMO7 is a novel hair cell-enriched protein, with specific localization to the cuticular plate.

LMO7 in other cell types was reported to localize at various subcellular sites. While most studies agree that LMO7 is localized at adherens junctions and focal adhesions, it was also reported that LMO7 might act as a nucleocytoplasmic shuttling protein to regulate transcription of emerin ^35^. In our immunolocalization studies, some antibodies showed weak nuclear staining (not shown), but the validity of that staining was deemed inconclusive. In order to ascertain the localization of endogenous LMO7 in hair cells, we employed the so-called split-GFP approach as an alternative method of protein localization (**Fig. 2a**). GFP features eleven beta strands making up a beta-barrel. Removing one of those beta-strands (creating the so-called GFP1-10) abolishes its fluorescence, but exogenous addition of the missing beta-strand (GFP11), which can be fused to a protein of interest, leads to spontaneous re-assembly and reconstitution of fluorescence. This strategy allows tagging and visualization of the protein of interest based on the localization of reconstituted GFP fluorescence. Split fluorescent protein has been widely used for protein quantification, visualization of protein cellular localization, as well as single-molecule imaging and protein-protein interaction^36-42^. Using CRISPR/Cas genome editing, we knocked-in the GFP11-coding sequence (48 nucleotides) into the C-terminus of mouse *Lmo7*, immediately prior to the stop codon (resulting in the *Lmo7-GFP11 KI mice)* (**Fig. 2b**). Explant cultures were established from *Lmo7-GFP11 KI* mice at P2, and to achieve complementation with GFP1-10, we transduced organ of Corti explants with AAV2/Anc80 virus carrying the GFP1-10 fragment coding sequence ^43^, driven by a CMV promoter (**Fig. 2c**). As illustrated in **Fig. 2d**, outer hair cells in *Lmo7-GFP11 KI*, which were transduced with GFP1-10, displayed robustly reconstituted GFP fluorescence at the cuticular plate level but not at the level of the cell body (**Fig. 2d,e**). Despite the fact that GFP1-10 was detected throughout the cytosol and the nucleus, we never detected any specific GFP fluorescence in the nucleus, suggesting that at least in hair cells, LMO7 is unlikely to regulate transcription by nucleocytoplasmic shuttling.

**Figure 2.**
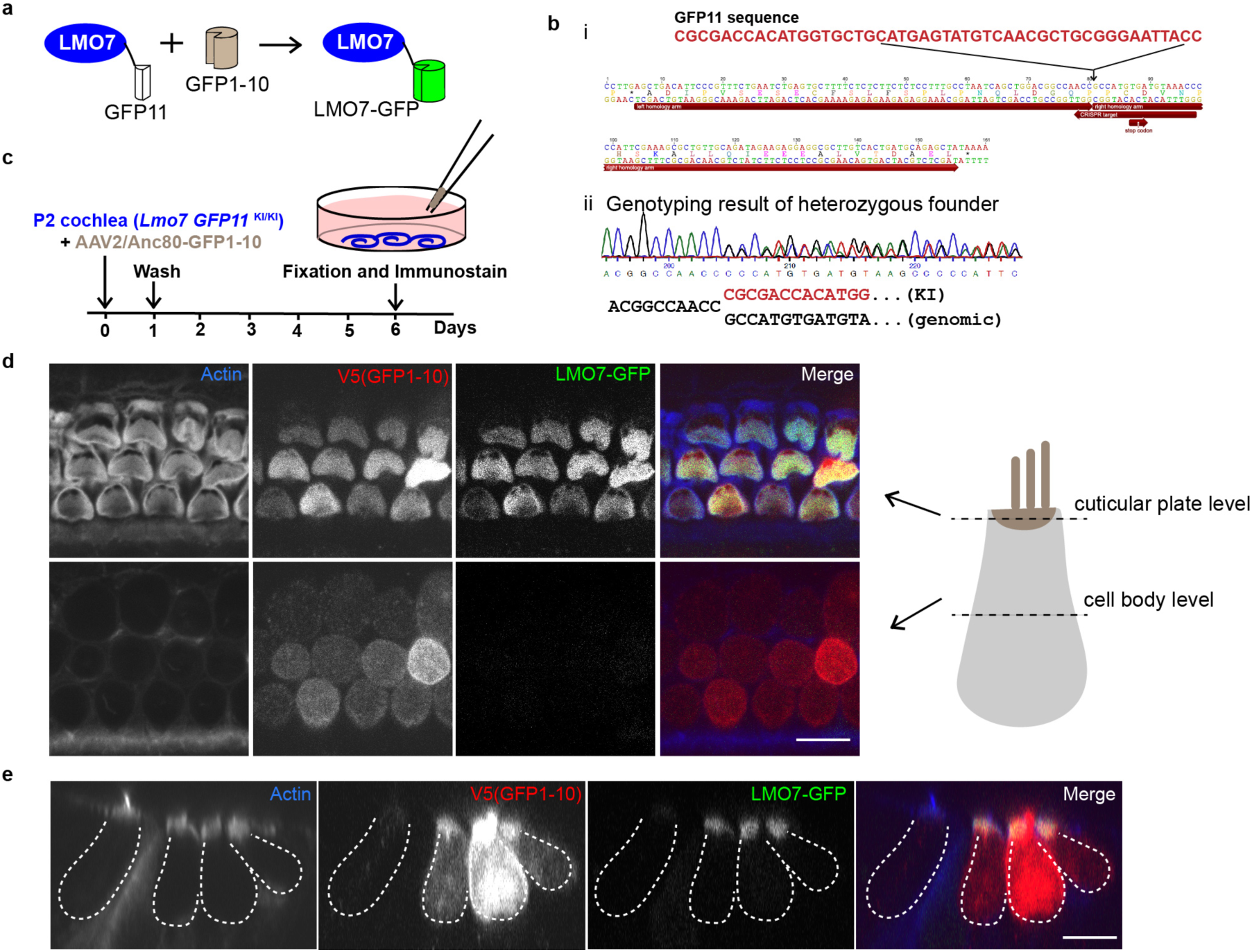
Localization of endogenous LMO7 using the split-GFP approach. **a**. Schematic diagram of the split-GFP approach to label tagged endogenous LMO7. **b**, Illustration of CRISPR/Cas9-mediated GFP11 knock-in strategy. (i) Genomic sequence surrounding the stop codon of mouse *Lmo7*. Highlighted are the CRISPR target sites and the left and right homology arms used for the homology-directed repair. (ii) Sequencing chromatogram of the modified locus, from the *Lmo7-GFP11 KI* founder mouse. **c.** Depiction of experimental paradigm for transduction of explants of *Lmo7-GFP11 KI* cochlea with Anc80-GFP1-10. The cochleae were dissected and transduced with Anc80-GFP1-10 for 24 hours. After replacement of culture medium, cultures were maintained for 5 days, fixed, and immunostained. **d**, Optical sections at the level of the cuticular plate and cell body are shown. Reconstituted LMO7-GFP expression localizes to the cuticular plate but not the cell body level. Scale bar, 5 μm. **e**, Side view of reconstituted LMO7-GFP expression enriched in the cuticular plate of outer hair cell. Scale bar, 5 μm.

### *Lmo7 KO* mice have defects in the cuticular plate

To investigate the function of LMO7 in the hair cell and in hearing function, we sought to analyze LMO7 loss-of-function mice. We initially obtained sperm of a mouse strain generated using the gene trap technology by Texas A&M ^44^ (mouse ID: Lmo7Gt(IST10208D3)Tigm) and generated the mice using *in vitro* fertilization. In these mice (*Lmo7 gene trap*), a cassette with a transcription-terminating polyadenylation site is inserted after exon 1. In a previous study, this mouse line was used to show that LMO7 is implicated in EDMD ^31^. However, immunolabeling experiments showed that LMO7 protein expression in hair cells of *Lmo7 gene trap* mice was not abolished (**Supplementary Fig. 1**). Immunoblot analysis further confirmed that the gene trap strategy only affected the longest isoform of LMO7 (**Fig. 3d**), consistent with the strategy to insert the gene trap cassette after exon 1. We therefore used CRISPR/Cas9 to generate a variety of mice with potentially deleterious mutations in the *Lmo7* gene. LMO7 is expressed in multiple, poorly characterized isoforms (**Fig. 3a**). In order to increase our chances of knocking out all isoforms, we performed three separate experiments to target exon 12, exon 17 and exon 28 of the mouse *Lmo7* gene. We isolated three founders, with a deleterious 2bp insertion in exon 12 (*Lmo7 exon12 KO* mouse), a 5bp deletion in exon 17 (*Lmo7 exon17 KO* mouse) or an 8bp deletion in exon 28 (*Lmo7 exon28 KO* mouse), respectively (**Fig. 3b and data not shown**). The founders were bred 4-5 generations, to breed out potential off-target mutations, and homozygotes were analyzed. In the *Lmo7 exon17 KO* mice, LMO7 immunoreactivity was undetectable, in auditory and vestibular hair cells (**Fig. 3c**), validating this mouse line as a *bona fide* loss-of-protein model. This was further confirmed by immunoblot analysis (**Fig. 3d**). The other mouse strains exhibited various degrees of incomplete ablation of LMO7 expression: *Lmo7 exon28 KO* mice showed reduced LMO7 immunoreactivity, and the LMO7 immunoreactivity is abolished in *Lmo7 exon12 KO* mice initially, but returns in the mature mice (**supplementary Fig. 1**). We believe that this is caused by a change in splicing, induced by the mutation. This phenomenon, while interesting, is not expected to be important for understanding LMO7’s function in hair cells. The remainder of the study thus focused on the *Lmo7exon17 KO* mouse line.

**Figure 3.**
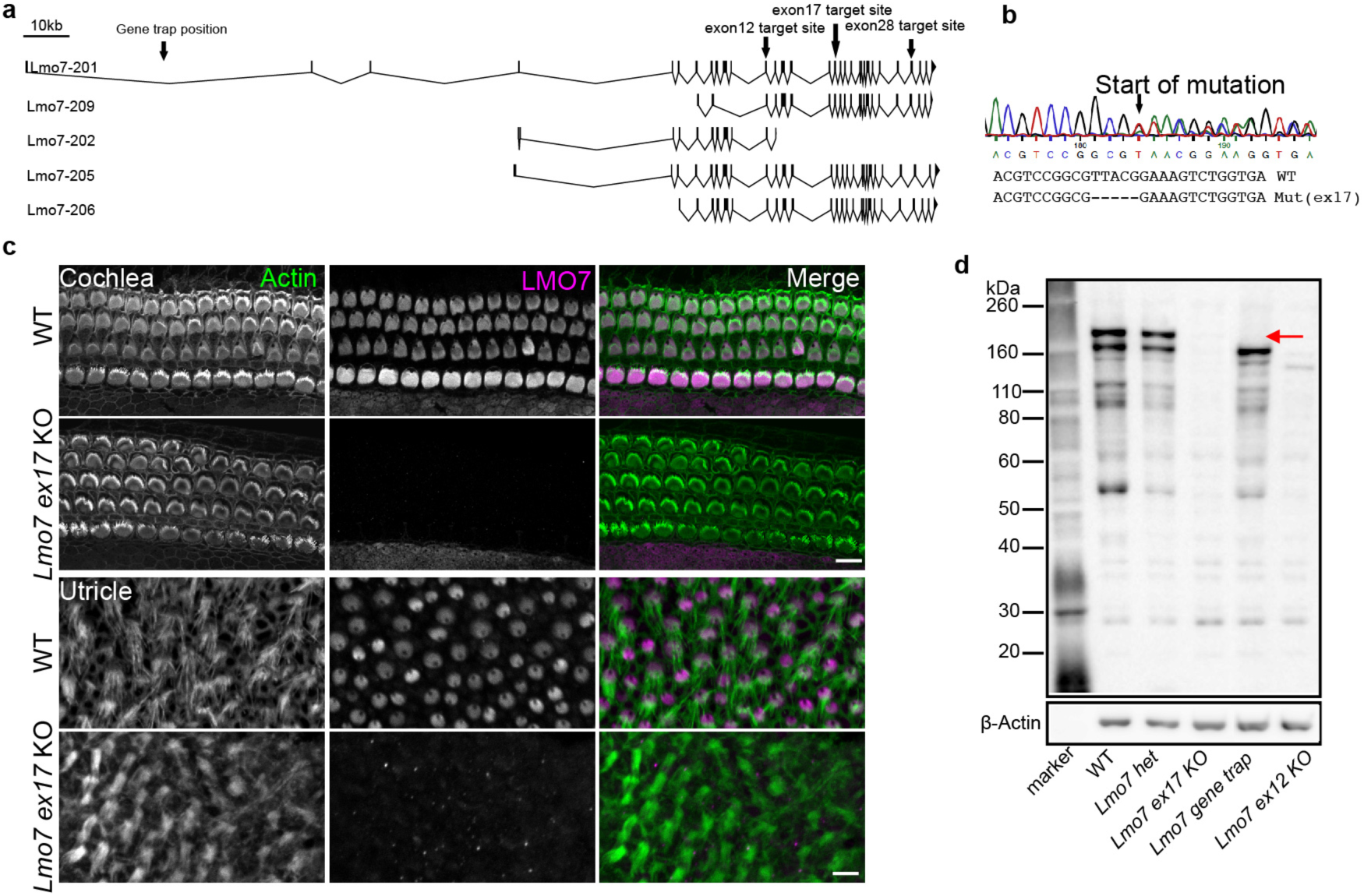
Generation of LMO7 deficient mice using CRISPR. **a**. Genomic structures of mouse *Lmo7* isoforms from Ensembl (GRCm38.p5). Exons and 5′ untranslated regions are indicated by black boxes, 3′ untranslated regions are indicated by black triangles. The locations of the gene trap and three CRISPR-mediated targeting sites are shown with arrows. **b.** Sanger sequencing result of edited locus demonstrates successful genome editing, resulting in a 5 bp deletion in exon 17. **c.** LMO7 immunolabeling in WT and *Lmo7 exon17 KO* mice in P4 cochlea and utricle. LMO7 is in magenta, and F-actin in green. LMO7 immunoreactivity is abolished in KO cochlear and vestibular hair cells. **d**, Comparative immunoblot of WT, *Lmo7 exon17 Het*, *Lmo7 exon17 KO, Lmo7 gene trap* and *Lmo7 exon12 KO* from P4 inner ear lysate, indicating loss of LMO7 expression in *Lmo7 exon17* and *Lmo7 exon17* lysate. The gene trap strategy only abolished the longest isoform, indicating by red arrow. The β-actin-specific bands indicate equal loading.

*Lmo7 exon17 KO* mice are healthy, viable, and display no overt phenotype. The gross morphology of the sensory epithelium was not affected. Upon closer inspection, however, some striking differences were detected: The cuticular plate of inner and outer hair cells showed a significantly reduced amount of F-actin, as measured by phalloidin staining (**Fig. 4a,b**). In outer hair cells, this phenotype progressed in an age-dependent manner (**Fig. 4b,c**). We next examined the rootlets that form the stereocilia insertion points into the cuticular plate. TRIOBP, an actin-bundling proteins, is an essential component of the rootlet, and TRIOBP deficiency leads to non-optimal mechanotransduction, causing hearing loss in mice and humans ^45^. We found that the spatial organization of the rootlets was disrupted in *Lmo7 exon17 KO* mice (**Fig. 4d,e**). In addition, the length of TRIOBP-positive rootlets, as well as the expression level of TRIOBP protein, was significantly reduced in the *Lmo7 exon17 KO* mice (**Fig. 4f**). In summary, LMO7 deficiency causes a reduction in F-actin density and disruption of the organization of the stereocilia rootlets in the cuticular plate.

**Figure 4.**
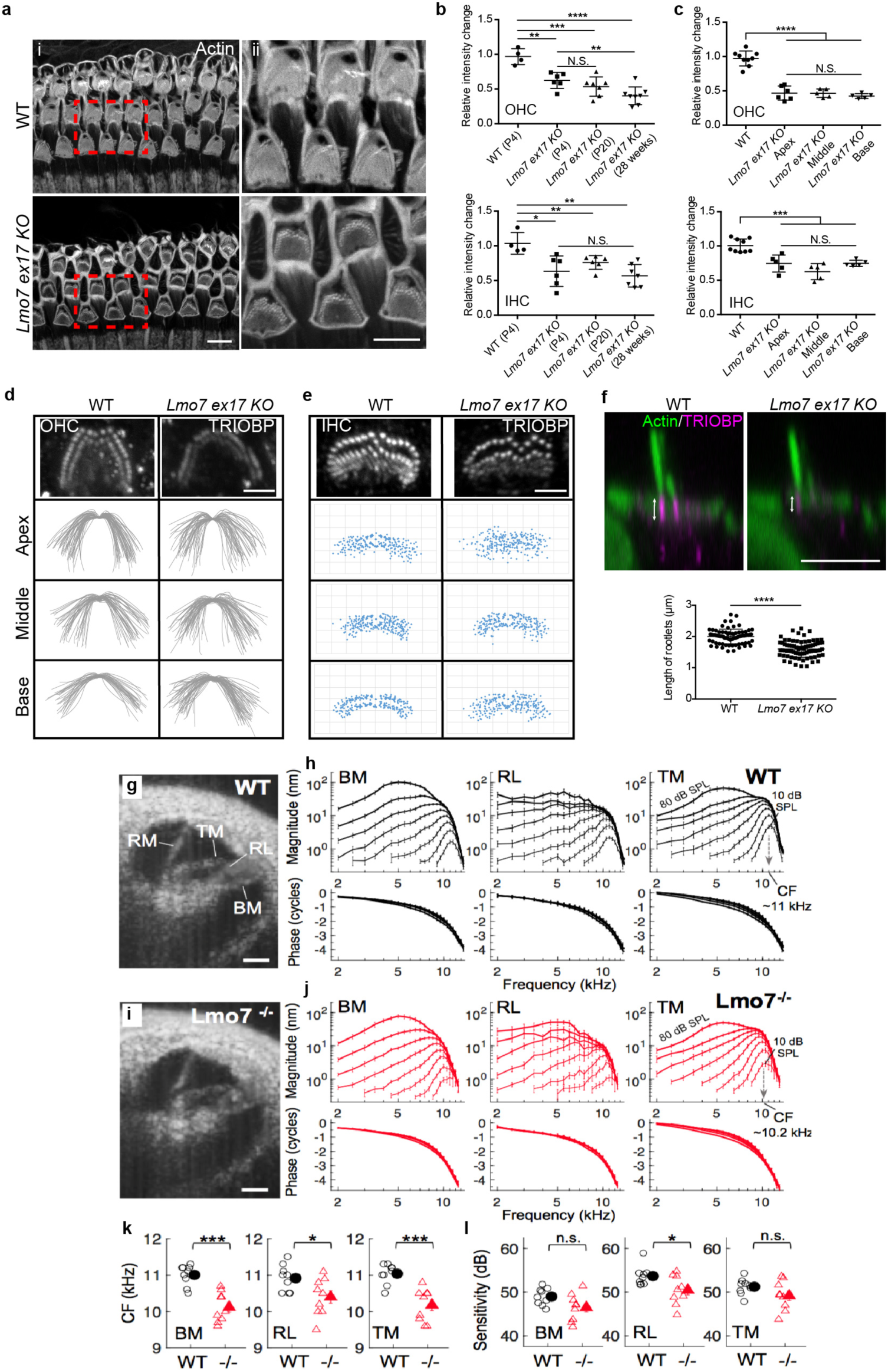
Reduced F-actin density and abnormal stereociliar rootlets in the cuticular plates of *Lmo7 exon17 KO* mice. **a.** (i) Reduced phalloidin staining in the cuticular plates of *Lmo7 exon17 KO* hair cells at P20. (ii) Enlarged view of boxed-in region. **b**. Quantification of relative grey value intensity changes of phalloidin reactivity in the cuticular plate of OHCs and IHCs between WT and *Lmo7 exon17 KO* mice, at different ages. The relative intensity changes are the ratio of phalloidin intensity of mutant and WT cuticular plates, at matched ages (n = 15-25 OHCs/mouse, N = 8 for P4 WT, N = 6 for P4 *Lmo7 exon17 KO* and WT, N = 7 for P20 *Lmo7 exon17 KO* and WT, N = 8 for 28 weeks *Lmo7 exon17 KO* and WT. n = 5-10 IHCs/mouse, N = 8 for P4 WT, N = 6 for P4 *Lmo7 exon17 KO* and WT, N = 6 for P20 *Lmo7 exon17 KO* and WT, N = 7 for 28 weeks *Lmo7 exon17 KO* and WT). **d.** Top view of TRIOBP staining in outer hair cell of WT and *Lmo7 exon17 KO* at p30. Lower three rows: line tracings of the first and second stereocilia rows in the apex, middle and base region., indicating a greater degree of variability in *Lmo7 exon17 KO* mice. **e.** Top view of TRIOBP staining in inner hair cells of WT and *Lmo7 exon17 KO* at p30. In inner hair cells, the rootlets were resolved well enough to allow scatter plot of individual rootlets of inner hair cells. n = 10 IHCs, N = 4 for WT and *Lmo7 ex17* KO. **f.** Side view of TRIOBP labeling in the rootlets at base of stereocilia of IHCs at P30. Below, quantitation of rootlet length of the longest row of stereocilia (as arrows shown in **f**) in inner hair cells at P30. N = 76 stereocilia, N = 4 for WT and *Lmo7 ex17 KO* Scale bar, 5 μm. Error bars indicate SD, ****p value<0.0001, ***p value<0.001, **p value<0.01, and *p value<0.05 in two-tailed unpaired Student’s t-test. **g. i.** Representative cross-sectional images of intact cochleae in live WT (**g**) and *Lmo7 exon17 KO* (**i**) mice obtained with VOCTV. Vibrations were measured from specific points on the basilar membrane (BM), reticular lamina (RL), and tectorial membrane (TM) in the apical cochlear turn. Reissner’s membrane (RM) is also indicated. Scale bars, 100 μm. **h,j,** Average sound-evoked displacement magnitudes and phases measured from the BM, RL, and TM in WT (**h**) and *Lmo7 exon17 KO* (**j**) mice (n = 9 in each group). Stimuli were swept from 2-14 kHz and varied in level from 10-80 dB SPL in 10 dB steps. TM displacement curves for the lowest and highest stimulus levels are labeled for clarity. All mice exhibited compressive, nonlinear growth of vibratory displacement magnitudes near the frequency evoking the largest displacements at low stimulus levels, i.e., the characteristic frequency (CF) of the recording location (downward-pointing arrows in b, d indicate the average CF). **k,** Average CFs for the BM, RL, and TM were significantly lower in *Lmo7 exon17 KO (-/-)* than in WT mice (individual data shown with open symbols). **l,** Vibratory sensitivity at CF also tended to be lower in *Lmo7 exon17 KO* mice than in WT, though only significantly so for the RL. Sensitivity was defined as the ratio of the response magnitude for a 20 dB vs. an 80 dB SPL tone at CF, after dividing both displacements by the respective stimulus pressure. Error bars indicate SEM. *p < 0.05, ***p < 0.0005, n.s. = p > 0.05, by unpaired t-test.

### LMO7 deficiency causes aberrant frequency tuning and reduced cochlear vibrations

We reasoned that the reduced F-actin density and aberrant rootlet organization in LMO7-deficient cuticular plates might reduce its stiffness. Such increase in compliance might impair the coupling of the reticular lamina and the tectorial membrane, affecting the sound-induced vibrations of cochlear partitions overall. We tested the vibratory movements of cochlear partitions *in vivo*, using volumetric optical coherence tomography and vibrometry (VOCTV) ^46^. The exquisite sensitivity of VOCTV enables the detection of subtle differences well before hearing threshold shift is evident. Therefore, to isolate the effect of cuticular plate defects on cochlear vibrations from the effects of age in general, and from the Cdh23(ahl) allele responsible for age-related hearing loss in the C57Bl6 strain, we performed VOCTV experiments at P30, on *Lmo7 exon 17* mice that were backcrossed to the CBA/J background. **Fig. 4g and i** show representative cross-sectional images of intact cochleae in live WT and *Lmo7 exon17 KO* mice obtained with VOCTV. Locations on the basilar membrane (BM), reticular lamina (RL), and tectorial membrane (TM) in the apical cochlear turn were selected for vibration measurements. Sound-evoked displacement magnitudes and phases measured from the BM, RL, and TM in WT and a *Lmo7 exon17 KO* mice are shown in **Fig. 4h and j**. Stimuli were swept from 2-14 kHz and varied in level from 10-80 dB SPL. Interestingly, the characteristic frequency (CF), which is defined as the frequency evoking the largest displacements at the lowest stimulus level (10 dB SPL), was slightly lower in the *Lmo7 ex17 KO* mouse (∽10 kHz) than in the WT mouse (∽11 kHz) for TM, as well as BM and RL (**Fig. 4k)**. A reduction in CF is consistent with the predicted reduction in stiffness in LMO7-deficient cuticular plates. Vibratory sensitivity at CF also tended to be lower in *Lmo7 exon17 KO* mice than in WT, though only significantly so for the RL (**Fig. 4l)**. Taken together, the VOCTV measurements indicate that LMO7 deficiency effects changes in the frequency tuning and sensitivity of cochlear vibrations.

### *Lmo7 exon17 KO* mice develop late-onset, progressive hearing loss, especially at low frequencies

Next, we examined the effects of the aforementioned defects of the cuticular plate and changes in cochlear vibrations on hearing function. We determined hearing thresholds in the *Lmo7 exon17 KO* mice, using auditory brainstem response (ABR) measurements. Despite the fact that abnormalities in the cuticular plate are detectable in early postnatal animals, and cochlear vibrations are altered at P30, hearing thresholds were comparable to WT controls up to 11 weeks of age (**Fig. 5a**). However, ABR thresholds became significantly elevated at 17 weeks, and worsened progressively, leading to near profound hearing loss at 26 weeks of age (**Fig. 5a**). Interestingly, while threshold differences were significant at most frequencies in the *Lmo7 exon17 KO* mice at 26 weeks of age, the lower frequencies were most affected. The hearing of *Lmo7 ex12 KO* mice were not affected until 26 weeks, while *Lmo7 gene trap* mice did not develop hearing loss at all (**Supplementary Fig. 2**). We also assessed OHC function using distortion product otoacoustic emissions (DPOAEs). At 17 weeks, a significant reduction in DPOAE levels was evident in the *Lmo7 ex17 KO* in mid-to low-frequency DPOAE output levels. DPOAE output was further reduced at 26 weeks of age (**Fig. 5b**).

**Figure 5.**
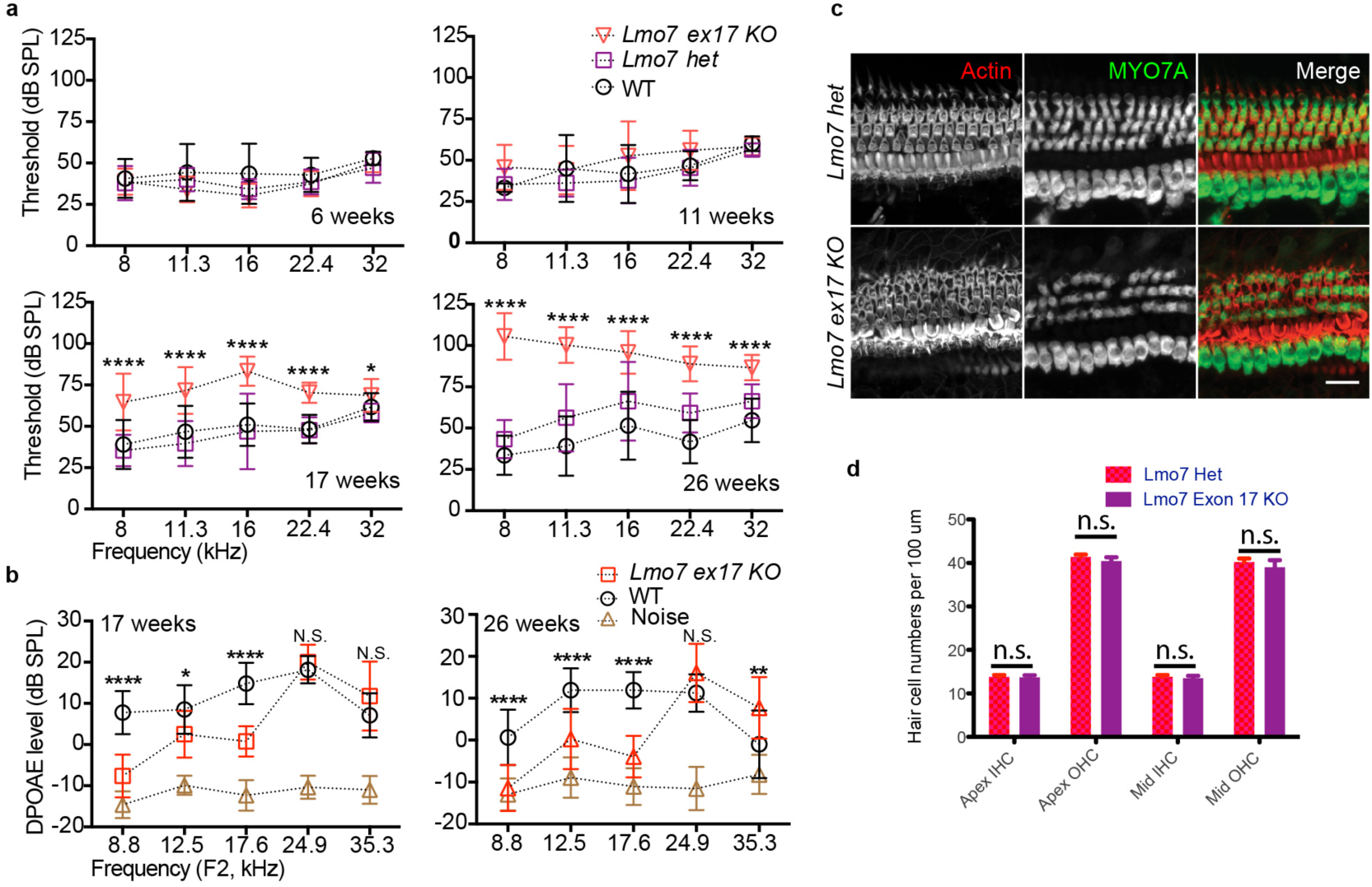
*Lmo7 exon17 KO* mice exhibit late onset, progressive hearing loss, especially at low frequencies, but do not lose hair cells. **a**. Auditory brainstem response (ABR) analysis demonstrates *Lmo7 exon17 KO* mice exhibit late onset, progressive hearing loss, especially at low frequencies. At 6 weeks: N = 7 for WT, N = 7 for *Lmo7* het, N = 16 for *Lmo7 exon17 KO*. At 11 weeks: N = 6 for WT, N = 13 for *Lmo7* het, N = 15 for *Lmo7 exon17 KO*. At 17 weeks: N = 15 for WT, N = 13 for *Lmo7* het, N = 15 for *Lmo7 exon17 KO*. At 26 weeks: N = 17 for WT, N = 12 for *Lmo7* het, N = 18 for *Lmo7 exon17 KO*. Both male and female mice were used. **b,** Distortion product otoacoustic emissions (DPOAEs) show significant differences in the frequency range (8-16 kHz) between *Lmo7 exon17 KO* and WT littermates at 17 and 26 weeks. At 17 weeks: N = 12 for WT, N = 13 for *Lmo7 exon17 KO*. At 26 weeks: N = 12 for WT, N = 14 for *Lmo7 exon17 KO*. Error bars indicate SD, ****p value<0.0001, **p value<0.01, and *p value<0.05 according to ANOVA test, followed by Tukey post-hoc analysis. **c,d,** Hair cells were counted based on MYO7A immunoreactivity over a length of 100 μm of the apical and middle turns of the cochlea. The hair cell numbers were not significantly affected in mutant mice. Number of animals *Lmo7 exon17 het* control: N = 5, for *Lmo7 exon17 KO:* N = 6. Scale bar, 10 μm.

To test whether hearing loss in *Lmo7 exon17 KO* mice is caused by loss of hair cells, we counted hair cell numbers in older mice after their terminal ABRs at 26 weeks. MYO7A immunoreactivity demonstrated that hair cell numbers were comparable between control and mutant cochleae (**Fig. 5c,d**), even in the apical regions corresponding to the frequencies that exhibit highly elevated thresholds in the ABR result of mutant mice. We conclude that *Lmo7 exon17 KO* mice develop late-onset, progressive hearing loss, especially at low frequencies. We further note that hearing loss is not caused by loss of hair cells.

### Stereocilia defects in older *Lmo7 exon17 KO* mice

We next investigated the cause for the progressive hearing loss observed in *Lmo7 exon17 KO* mice that especially impaired low-frequency hearing. The reduced DPOAE output starting at ∽17 weeks of age could have its origin in changes in cochlear vibration properties, as documented by VOCTV measurements. The reduced vibrational sensitivity however is unlikely to be the sole cause for the near profound hearing loss at 26 weeks of age. To identify other pathological correlates for functional hearing loss, we examined the ultrastructure of hair cells in *Lmo7 exon17 KO* mice, using scanning electron microscopy (SEM). At 11 weeks of age, the morphology of hair cells in *Lmo7 exon17 KO* is comparable to WT controls (**Fig. 6a,b**). The reduced F-actin density evident in phalloidin immunofluorescence thus does not cause overt changes in the surface morphology of the cuticular plate. Some hair bundles of outer hair cells of *Lmo7 exon17 KO* mice display asymmetry of their characteristic W-shape (**Fig. 6a**), consistent with the aberrant TRIOBP staining at younger ages. At 26 weeks, the outer hair cell morphologies were comparable in appearance between WT and *Lmo7 exon17 KO* mice. In contrast, profound defects in inner hair cell stereocilia were detected, such as stereocilia fusions and loss. This phenotype was restricted to the apical cochlear regions (**Fig. 6d,e**). More subtle changes in inner hair cell stereocilia were also evident in the other regions of the cochlea: the “tenting” of the second row of stereocilia, thought to be a correlate for functional tip links and active mechanotransduction (MET), was reduced in inner hair cell stereocilia of mid-basal turns *Lmo7 exon17 KO* mice (**Fig. 6d,f**).

**Figure 6.**
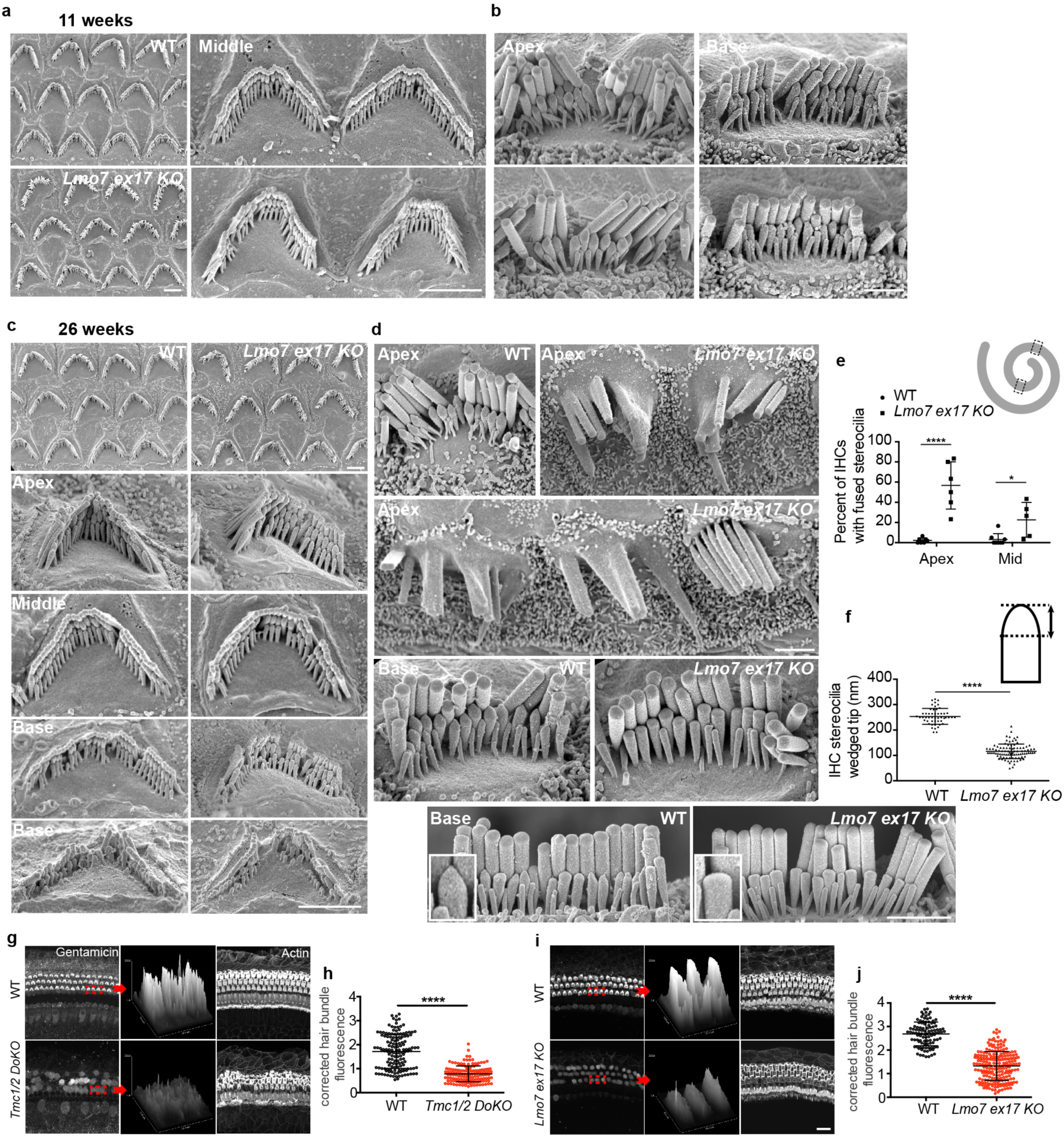
Stereocilia defects in older *Lmo7 exon17 KO* mice. **a.** Scanning electron micrographs (SEMs) of outer hair cells and **b.** inner hair cells show no overt abnormalities at 11 weeks of age. **c.** SEMs of outer hair cells show comparable hair bundle morphology in WT and *Lmo7 exon17 KO* mice at 26 weeks of age. Both genotypes show degeneration of basal hair cell stereocilia, which is expected in the BL6 background. **d.** Inner hair cells in the apical region of *Lmo7 exon17 KO* have high incidence of fused stereocilia. Fusions were not detected in WT controls. In basal hair cells of *Lmo7 exon17 KO* mice, stereocilia tenting was significantly reduced compared to WT controls, indicating reduced mechanotransduction activity. **e.** Quantification of percent of IHCs with fused stereocilia within the regions labeled with dotted boxes (upper right cartoon), (n = 5 cochleae for WT Apex, n = 6 cochleae for Lmo7 ex17 KO Apex, n = 6 cochleae for WT Middle, n = 5 cochleae for Lmo7 ex17 KO Middle, N = 4 for both groups). **f**. Heights of the basal IHC stereocilia wedged tips (upper right cartoon) in the second row of the hair bundle (n = 50 stereocilia from 9 WT cells, n = 92 stereocilia from 13 Lmo7 exon17 KO cells, N = 4 for both groups). Error bars indicate SD, ****p value<0.0001 and *p value<0.05 in two-tailed unpaired Student’s t-test. **g.** Validation of gentamicin uptake assay in *Tmc1/2 DOKO* mice at P28. Scale bar, 10 μm. **h,** Quantitation of corrected hair bundle gentamicin uptake in WT and *Tmc1/2 DOKO mice*, n = 148 OHCs, N = 3 for WT, n = 144 OHCs, N = 4 for *Tmc1/2 DOKO*. **i,** Gentamicin immunoreactivity, especially in the hair bundles, is significantly reduced in *Lmo7 exon17 KO* mice at 28 weeks, compared to WT controls. Scale bar, 10 μm. **j,** Quantitation of corrected hair bundle gentamicin uptake in WT and *Lmo7 exon17 KO* mice. n = 104 OHCs, N = 3 for WT, n = 214 OHCs, N = 3 for Lmo7 ex17 KO mice. Error bars indicate SD, ****p value<0.0001 in two-tailed unpaired Student’s t-test.

We next examined whether the stereocilia abnormalities could affect hair cell MET in 26 week old *Lmo7 exon17 KO* mice. In the absence of an established method for assessing hair cell MET function in adult mice on a cellular level, we applied and quantified the uptake of the aminoglycoside gentamicin into hair cells *in vivo* as a surrogate for MET activity. Gentamicin was given i.p. in conjunction with the loop diuretic furosemide which increases permeation of the blood-labyrinth-barrier ^47,48^. As proof of principle, we first validated the usefulness of this technique as a readout for MET function, using MET-deficient *Tmc1/2 double KO (DOKO)* as negative controls. Mice were injected at P28, and were sacrificed 2 hours later, prior to the onset of gentamicin-mediated ototoxicity. Cochleae were processed for staining with a gentamicin-specific antibody. As described in **Fig. 6g and h**, hair cells in WT mice at P28 displayed robust uptake of gentamicin, especially in the stereocilia. Gentamicin immunoreactivity was stronger in outer hair cells, similar to the pattern of FM1-43 uptake we observe in neonatal explants (data not shown). Gentamicin immunoreactivity in *Tmc1/2 DOKO* hair cells was significantly reduced, and was not concentrated in the stereocilia as in WT controls (**Fig. 6g and h)**. Some remaining gentamicin immunoreactivity was detected in the *Tmc1/2 DOKO* hair cells, likely because gentamicin enters hair cells through routes other than the MET channel, including endocytosis ^49,50^. Having validated this method, we applied this method to tested MET activity at 28 weeks, when *Lmo7 exon17 KO* mice exhibit near profound hearing loss. Compared to WT mice, *Lmo7 exon17 KO* mice showed a clear difference in gentamicin immunoreactivity, in both inner and outer hair cells (**Fig. 6i,j**). We therefore conclude that the hearing loss in *Lmo7 ex17 KO* mice correlates with a reduction in hair cell function as inferred from gentamicin uptake.

## Discussion

While the impact of various genetic and environmental stressors on hair bundle integrity and hair cell function is well studied, the effect of cuticular plate defects on mammalian hearing function is not well understood. Our investigations into the role of LMO7, which we show is a protein specifically localized to the cuticular plate, provides a unique avenue into a better understanding of the significance of the cuticular plate for the performance of the hair cell and the cochlea.

Our findings are consistent with a role of LMO7 in mediating actin polymerization in the cuticular plate. This is in agreement with a previous study reporting that LMO7 in Hela cells increased the ratio of filamentous to globular actin (F/G-actin), resulting in the activation of the transcription factor serum response factor (SRF) ^32^. The precise mechanism by which LMO7 influences actin polymerization is not known, but LMO7 was reported to act as a scaffolding protein, providing a platform for Cdc42 and Rac1 activity, thereby influencing actin dynamics towards more actin polymerization ^32^. Furthermore, LMO7 is known to bind alpha-actinin, an actin-crosslinker resident in the cuticular plate ^51^. LMO7 also harbors a calponin homology, PDZ and LIM domain, all of which are promiscuous protein interaction modules ^28,29^. We posit that LMO7 is a factor that promotes the polymerization and crosslinking of the F-actin cytoskeleton in the cuticular plate.

The primary effect resulting from LMO7 deficiency is a reduction of F-actin density and abnormal rootlet organization in the cuticular plate, evident at early postnatal stages. We propose that the expected reduction of the cuticular plate stiffness affects cochlear mechanics, specifically causing reduced tuning and sensitivity of cochlear vibrations, as shown in the VOCTV measurements. The cuticular plate defects in *Lmo7 exon17 KO* mice are initially well compensated, but we propose that progressive degradation of the cuticular plates and cochlear vibrations ultimately affect DPOAE output and hearing thresholds starting at around 17 weeks of age.

In addition to the direct and early effects on the cytoskeletal organization of the cuticular plate, LMO7 deficiency causes a late-onset degeneration of the inner hair cell stereocilia. In the apical and middle regions of 26 weeks old mutant mice, many stereocilia were fused, and in the base, the transducing stereocilia of the second row displayed significantly less tenting, indicative of reduced mechanotransduction. This was consistent with a reduction of gentamicin uptake, which we used as a crude and indirect measure for hair cell mechanotransduction activity. We propose that the stereocilia degeneration is the cause for the profound hearing loss in older mice, especially affecting low frequencies. Since LMO7 is not localized in the stereocilia, we suggest that the degeneration of stereocilia occurs as a result of the cuticular plate defects. One could speculate that fusion and tenting defect could be a result of decreased cuticular plate rigidity and lack of rootlet organization, which allows stereocilia to touch and fuse.

Alternatiavely, LMO7 deficiency might disrupt the G/F-actin homeostasis and SRF signaling in hair cells, as previously shown in HeLa cells ^32^. Repression of SRF signaling, considered a master regulator of actin homeostasis ^52-56^, could affect the F-actin cytoskeleton in the entire hair cell, including the F-actin-based stereocilia. Interestingly, hair cell specific deletion of *Cdc42*, a member of the Rho-family of small GTPases, and a well-established upstream factor of SRF activity, causes stereocilia fusions and depletion^57^.

It was proposed that LMO7 is a nuclear shuttling protein ^35,58^, with the potential to relay information about the tensional state of the epithelium into transcriptional activity in the nucleus. In our analysis, neither antibody-based nor direct localization of tagged endogenous LMO7 using the split-GFP approach provided any convincing localization of LMO7 in the nucleus. At least in hair cells, it is thus unlikely that LMO7 plays a role in nuclear transcriptional activity.

Most genetic or environmental ototoxic stressors primarily impact high-frequency hearing, with the low frequency hearing affected to a lesser degree and with a delayed onset. Accordingly, most genetic models for hearing loss focus on high frequency hearing loss. The *Lmo7 exon 17 KO* mouse is a rare experimental model for low frequency hearing loss, providing an opportunity to study the molecular mechanisms underlying this understudied type of hearing loss.

## Methods

### Animal care and handling

The protocol for care and use of animals was approved by the University of Virginia Animal Care and Use Committee. The University of Virginia is accredited by the American Association for the Accreditation of Laboratory Animal Care. C57BL/6 (Bl6) and CBA/J mice used in this study were ordered from Jackson Laboratory (ME, USA). All mouse experiments were performed using C57BL/6 or CBA/J (Cochlear vibration measurements) inbred mouse strains. Neonatal mouse pups [postnatal day 0 (P0)-P4] were killed by rapid decapitation, and mature mice were killed by CO2 asphyxiation followed by cervical dislocation. *Lmo7 gene trap* mice were obtained from Dr. James M. Holaska (University of Chicago). *Tmc1/2 DOKO* mice were provided by Dr. Andrew Griffith. *Lmo7 exon17 KO* mice were bred ten generations to the C57BL/6 or CBA/J.

### Immunocytochemistry

Tissues were fixed for 30 minutes in 3% formaldehyde. After blocking for 1 h with 1% bovine serum albumin, 3% normal donkey serum, and 0.2% saponin in PBS (blocking buffer), organs were incubated overnight at 4°C with primary antibodies in blocking buffer. Organs were then washed 5 min with PBS and incubated with secondary antibodies (7.5 μg/ml Alexa 647, Alexa 488, Alexa 555-conjugated donkey anti-rabbit IgG, donkey anti-mouse IgG, donkey anti-goat IgG, Invitrogen,) and 0.25 μM phalloidin-Alexa 488 (Invitrogen) in blocking buffer for 1-3 h. Finally, organs were washed five times in PBS and mounted in Vectashield (Vector Laboratories). Samples were imaged using Zeiss LSM880 and Leica confocal microscopes. The following antibodies were used: rabbit anti-LMO7 (M-300, sc-98422; Santa Cruz Biotechnology), mouse anti-Spectrin alpha chain (MAB1622; MilliporeSigma), V5 Tag Antibody (R960-25, Invitrogen), rabbit anti-TRIOBP (16124-1-AP, Proteintech Group), rabbit anti-MYO7A (111501, Proteus BioSciences), mouse anti-Gentamicin (16102, QED Bioscience Inc.)

### Immunoblots

The whole P4 inner ear were homogenized and protein were isolated using Nucleic Acid and Protein Purification kit (Macherey-Nagel). Half of an inner ear per well was loaded onto the 12% Bis-Tris SDS PAGE gel (Novex 4–12%, Invitrogen), transferred to PVDF membranes, and stained with India Ink. Blots were then blocked in blocking buffer (ECL prime blocking reagent; GE Healthcare) for 1 h and probed with primary antibodies overnight at 4°C. After three 5 min washings in TBS/0.1% Tween 20, blots were incubated with HRP-conjugated goat anti-rabbit secondary antibody (Cell Signaling Technology) for 1 h, and bands were visualized by ECL reagent (Pierce Biotechnology ECL Western blotting substrate and GE Healthcare GE ECL prime Western blotting reagent). Chemiluminescence was detected using an ImageQuant LAS4000 mini imager (GE Healthcare).

### CRISPR/Cas-mediated generation of *Lmo7 KO mice* and *Lmo7 GFP11 KI mice*

Cas9 endonuclease mRNA was generated using the plasmid MLM3613 (provided by Keith Joung’s lab through Addgene) as a template. The target sequence was chosen using the “CRISPR Design” bioinformatics tool, developed by Feng Zhang’s lab at the Massachusetts Institute of Technology (crispr.mit.edu). The targets sequences used were as follows: CCTGCGTCAGGTGCGCTACG on Lmo7 exon 12, GTGATGGACGTCCGGCGTTA on Lmo7 exon 17, AGAGGATGGGTTCCGCATGT on Lmo7 exon 28, GGTTTACATCACATGGCGGT on Lmo7 exon 31 for GFP11 KI) were cloned downstream of the T7 promoter in the pX330 vector (provided by Feng Zhang’s lab through Addgene). IVT was performed using the MAXIscript T7 kit (Life Technologies) and RNA was purified using the MEGAclear kit (Life Technologies). The repair template (GAGCTGACATTCCCGTTTCTGAATCTGAGTGCTTTTCTCTCTTCTCTCCTTTGCCTAATCAGCTGGACGGCCAACCCGCGACCACATGGTGCTGCATGAGTATGTCAACGCTGCGGGAATTACCGCCATGTGATGTAAACCCCCATTCGAAAGCGCTGTTGCAGATAGAAGAGGAGGCGCTTGTCACTGATGCAGAGCTA) for GFP11 KI was made by Integrated DNA Technologies (Ultramer, PAGE-purified). The *Lmo7 KO* mice and *Lmo7 GFP11 KI* mice were generated according to the procedure published previously ^59^. Briefly, fertilized eggs produced from B6SJLF1 (The Jackson Laboratory) mating were co-injected with Cas9 protein (PNA Bio, 50 ng/μl) and sgRNA (30 ng/μl) with or without GFP11 repair template (10 ng/ul). Two-cell stage embryos were implanted on the following day into the oviducts of pseudopregnant ICR female mice (Envigo). Pups were screened by PCR and founders identified by DNA sequencing of the amplicons for the presence of indels and/or repair.

### AAV vector and organotypic explant cultures

The scCMV-GFP1-10-W3SL plasmid cassette was made by Dr. Edward Perez-Reyes (University of Virginia). AAV2/Anc80-GFP1-10 vector was obtained from Gene Transfer Vector Core at Schepens Eye Research Institute, Massachusetts Eye and Ear. Titer of AAV stock 4 ×; 10^12^ GCs/mL. Mouse cochleae were dissected in Hank’s balanced salt solution (HBSS, Invitrogen, MA) containing 25 mM HEPES, pH 7.5. The organ of Corti was separated from the spiral lamina and the spiral ligament using fine forceps and attached to the bottom of sterile 35 mm Petri dishes (BD Falcon, NY), with the hair bundle side facing up. The dissection medium was then replaced by two exchanges with culture medium (complete high-glucose DMEM containing 1% FBS, supplemented with ampicillin and ciprofloxacin). Prior to experimental manipulation, explants were pre-cultured for 24h, to allow acclimatization to the culture conditions (Francis et al. 2013). The cochleae were transduced with AAV2/Anc80-GFP1-10 for 24 hours (virus was diluted 1:200 in the culture medium). After replacement of culture medium, cultures were maintained for 5 days, then fixed, and immunostained. The GFP1-10 coding sequence was followed by a DNA sequence coding for the V5 affinity tag. V5 immunoreactivity hence serves as transfection control.

### Auditory brainstem response

Auditory brainstem responses (ABRs) of adult WT, heterozygous and homozygous *Lmo7 exon 17 KO*, *Lmo7 exon 12 KO*, *Lmo7 gene trap KO* mice were recorded from ages 6 to 26 weeks. Mice were anesthetized with a single intraperitoneal injection of 100 mg/kg ketamine hydrochloride (Fort Dodge Animal Health) and 10 mg/kg xylazine hydrochloride (Lloyd Laboratories). All ABRs were performed in a sound-attenuating booth (Med-Associates), and mice were kept on a Deltaphase isothermal heating pad (Braintree Scientific) to maintain body temperature. ABR recording equipment was purchased from Intelligent Hearing Systems. Recordings were captured by subdermal needle electrodes (FE-7; Grass Technologies). The noninverting electrode was placed at the vertex of the midline, the inverting electrode over the mastoid of the right ear, and the ground electrode on the upper thigh. Stimulus tones (pure tones) were presented at a rate of 21.1/s through a high-frequency transducer (Intelligent Hearing Systems). Responses were filtered at 300-3000 Hz and threshold levels were determined from 1024 stimulus presentations at 8, 11.3, 16, 22.4, and 32 kHz. Stimulus intensity was decreased in 5-10 dB steps until a response waveform could no longer be identified. Stimulus intensity was then increased in 5 dB steps until a waveform could again be identified. If a waveform could not be identified at the maximum output of the transducer, a value of 5 dB was added to the maximum output as the threshold.

### Distortion product otoacoustic emissions

Distortion product otoacoustic emissions (DPOAE) of adult WT and homozygous Lmo7 ex17 KO mice were recorded at 17 and 26 weeks. While under anesthesia for ABRs testing, DPOAE were recorded firstly using SmartOAE ver. 5.20 (Intelligent Hearing Systems). A range of pure tones from 8 to 32 kHz (16 sweeps) was used to obtain the DPOAE for right ear. DPOAE recordings were made for *f*_2_ frequencies from 8.8 to 35.3 kHz using paradigm set as follows: *L*_1_ = 65 dB, *L*_2_ = 55 dB SPL, and *f*_1_/*f*_2_ = 1.22.

### Hair cell counts

Mice were killed after the final ABRs at 26 weeks. Cochleae were dissected, openings were created at the base and apex of the cochlea and fixed in 4% paraformaldehyde (PFA) (RT-15720, Electron Microscopy Science, PA) for 24h. After decalcification for 7 days in EDTA solution, apical and middle turns of the cochlear sensory epithelium were dissected. Hair cell counting was performed for the organ of Corti using MYO7A immunoreactivity as a marker of hair cell presence. After confocal microscopy, images were analyzed using ImageJ. Hair cells were counted from the apical (0.5-1 mm from apex tip, corresponding to the 6-8 kHz region), mid turns (1.9-3.3 mm from the apex tip, corresponding to 12-24 kHz region) and of the cochlea. Organ of Corti from at least 5 mice were analyzed for each experimental condition.

### Scanning electron microscopy

Adult mice were killed via CO2 asphyxiation; animals were perfused intracardially, with 2.5% glutaraldehyde + 2% formaldehyde. The cochlea was dissected and a piece of bone was removed from the apex to create an opening. The stapes and oval window were then removed, and fixative (2.5% glutaraldehyde, in 0.1 M cacodylate buffer, with 3 mM CaCl2) was perfused through the apical and basal openings. Cochleae were incubated in fixative overnight at 4°C, then were decalcified in 4.13% EDTA, pH 7.3, for 10 days at room temperature. After the tectorial membrane was removed, the organ of Corti was dissected from the cochlea and was post-fixed in 1% OsO4, 0.1 M cacodylate buffer, and 3 mM CaCl2. The tissue was then processed according to the thiocarbohydrazide-OsO4 protocol (Davies and Forge, 1987). Cochleae were then dehydrated through a series of graded ethanol incubations, critical point dried, and mounted on stubs. After sputter coating with gold, they were imaged on a Zeiss Sigma VP HD field emission SEM using the secondary electron detector.

### Cochlear vibration measurements

Volumetric optical coherence tomography and vibrometry (VOCTV) was used to image and measure vibrations from the intact mouse cochlea, as described in Lee et al. (2015, 2016). Mice (P28-P32) of either sex were anesthetized with ketamine/xylazine (80-100mg/kg), placed on a heating pad, and the skull was fixed to a custom head-holder with dental cement. A ventrolateral surgical approach was then used to access the left middle ear bulla, which was widely opened so that the otic capsule bone and middle ear ossicles could be visualized. VOCTV was performed using a custom-built system, which consisted of a broadband swept-source (MEMS-VCSEL, Thorlabs) with a 1310 nm center wavelength, 100 nm bandwidth, and 200 kHz sweep rate, as well as a dual-balanced photodetector (WL-BPD600MA, Wieserlabs), and a high-speed digitizer (NI-5761, National Instruments). The source beam was scanned across the preparation using a 2-D galvo mirror housed in an adaptor attached to the bottom of the dissecting microscope (Stemi-2000, Zeiss) to obtain cross-sectional images of the apical cochlear turn. Sound-evoked vibrations were then measured from specific points on the basilar membrane (BM), reticular lamina (RL) and tectorial membrane (TM), with 100 ms pure-tones presented via a speaker (MDR EX37B, Sony) positioned close to the eardrum. Displacement magnitudes and phases were obtained with stimulus frequency ranging from 2-14 kHz in 0.5 kHz steps and stimulus level varied from 10-80 dB SPL in 10 dB steps. After sacrificing the mouse via anesthetic overdose, displacements of the middle ear ossicular chain were measured. This was done so that the middle ear response phase could be subtracted from the cochlear vibration phase at each frequency, thus eliminating the influence of middle ear transmission delays. All displacement responses analyzed and shown here were required to have magnitudes falling at least 3 standard deviations above the mean of the measurement noise floor at surrounding frequencies.

### Gentamicin uptake experiment

Gentamicin sulfate (Fisher Scientific), was dissolved in 0.9% sterile saline to a concentration of 50 mg/ml. Adult mice received one subcutaneous injection of 100 mg/kg gentamicin. This was followed by a single intraperitoneal injection of furosemide (10 mg/ml; Hospira) at a dosage of 400 mg/kg 30 min later. The furosemide injection was used to enhance gentamicin uptake by increasing permeation of the blood-labyrinth-barrier. Mice were sacrificed 2 hours later, prior to the onset of gentamicin-mediated ototoxicity. Cochleae were processed for staining with a gentamicin-specific antibody.

### Statistics

For statistical analysis, GraphPad Prism (La Jolla, CA) was used. Two-way analysis of variance (ANOVA) was used to determine statistically significant differences in the ABR and DPOAE analyses. Significant differences in individual frequencies were determined by a Tukey post-hoc analysis. For two-tailed unpaired Student’s t-test, P-values smaller than 0.05 were considered statistically significant. All n in statistical analyses refer to numbers of stereocilia, hair cells or cochlea regions, N in statistical analyses refer to number animals. All error bars indicate standard deviation (SD) or standard error of the mean (SEM).

## Supplementary Figures

**Supplementary Figure 1.**
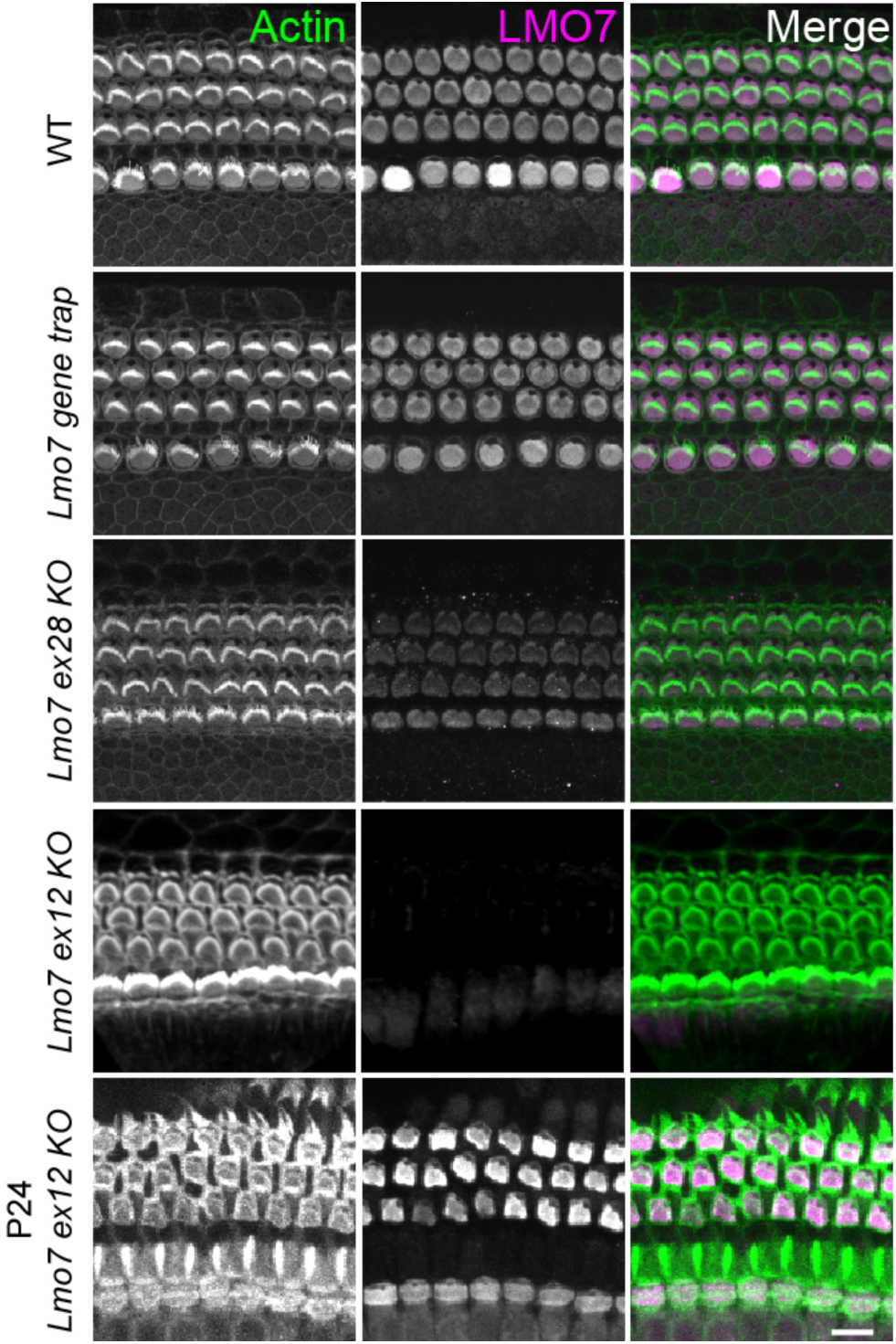
LMO7 expression pattern in different mutant mice. Immunocytochemistry for LMO7 expression in P4 WT, *Lmo7 gene trap*, *Lmo7 ex28 KO*, *Lmo7 ex12 KO* and P24 *Lmo7 ex12 KO* mice cochlea.

**Supplementary Figure 2.**
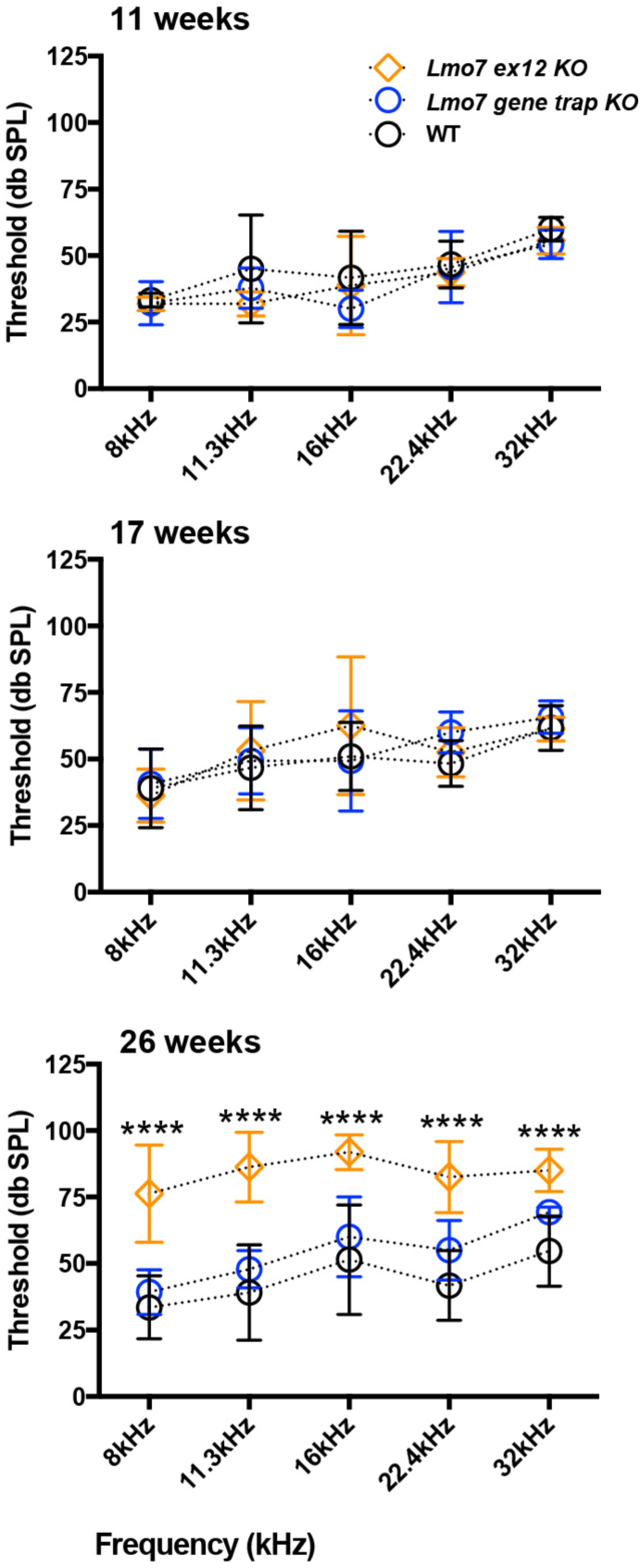
Auditory brainstem response analysis demonstrates that *Lmo7 exon12 KO* but not *Lmo7 gene trap* mice develop late onset, progressive hearing loss. At 11 weeks: N = 6 for WT, N = 7 for *Lmo7 gene trap*, N = 8 for *Lmo7 exon12 KO*. At 17 weeks: N = 15 for WT, N = 7 for *Lmo7 gene trap*, N = 8 for *Lmo7 exon12 KO*. At 26 weeks: N = 17 for WT, N = 7 for *Lmo7 gene trap*, N = 8 for *Lmo7 exon12 KO*. Both male and female mice were used. Error bars indicate SD, ****p value<0.0001 according to ANOVA test, followed by Tukey post-hoc analysis.

**Author contributions**
TTD, JBD, EW, SP and JBS performed the experiments. TTD, JBD and JBS analyzed the data. EPR, WX and JSO provided material and support. TTD and JBS wrote the manuscript.

## Acknowledgements

We thank the Virginia Lion’s Hearing Foundation (VLHF) for the generous financial support. This study was supported by NIH R01, 1R01DC014254.

